# Palmitoylation of the p75 neurotrophin receptor is required for retrograde regressive signaling

**DOI:** 10.64898/2026.10.02.756216

**Authors:** Amrita Pathak, Hrishita Das, Pooja Shukla, Austin B. Keeler, Christopher D. Deppmann, Bruce D. Carter

**Author notes:** equal contributions (Christopher D. Deppmann, Bruce D. Carter).

## Abstract

Target-derived neurotrophins regulate neuronal survival and pruning via retrograde signaling. Unlike Trk receptors, which are transported as transmembrane proteins, the p75 neurotrophin receptor (p75NTR) undergoes regulated intramembrane proteolysis to release a soluble intracellular domain (ICD) that must somehow associate with axonal endosomes for retrograde transport. Here, we identify palmitoylation of a conserved juxta membrane cysteine (C281) as a critical requirement for this process. In sympathetic neurons, we demonstrate that p75NTR palmitoylation is dispensable for surface trafficking and proteolytic cleavage but is strictly required for ligand-induced internalization and the biogenesis of p75ICD-positive signaling endosomes. Consequently, blocking palmitoylation, either pharmacologically or via a palmitoylation-deficient C281A mutation, prevents the accumulation of p75ICD in axonal endosomes and blocks axon-initiated apoptosis. *In vivo*, C281A knock-in mice exhibit reduced developmental apoptosis of sympathetic neurons, similar to p75NTR deletion. These findings define palmitoylation as an essential checkpoint that couples p75NTR cleavage to endosomal segregation, thereby enabling the long-distance transmission of regressive signals.

**Synopsis:** 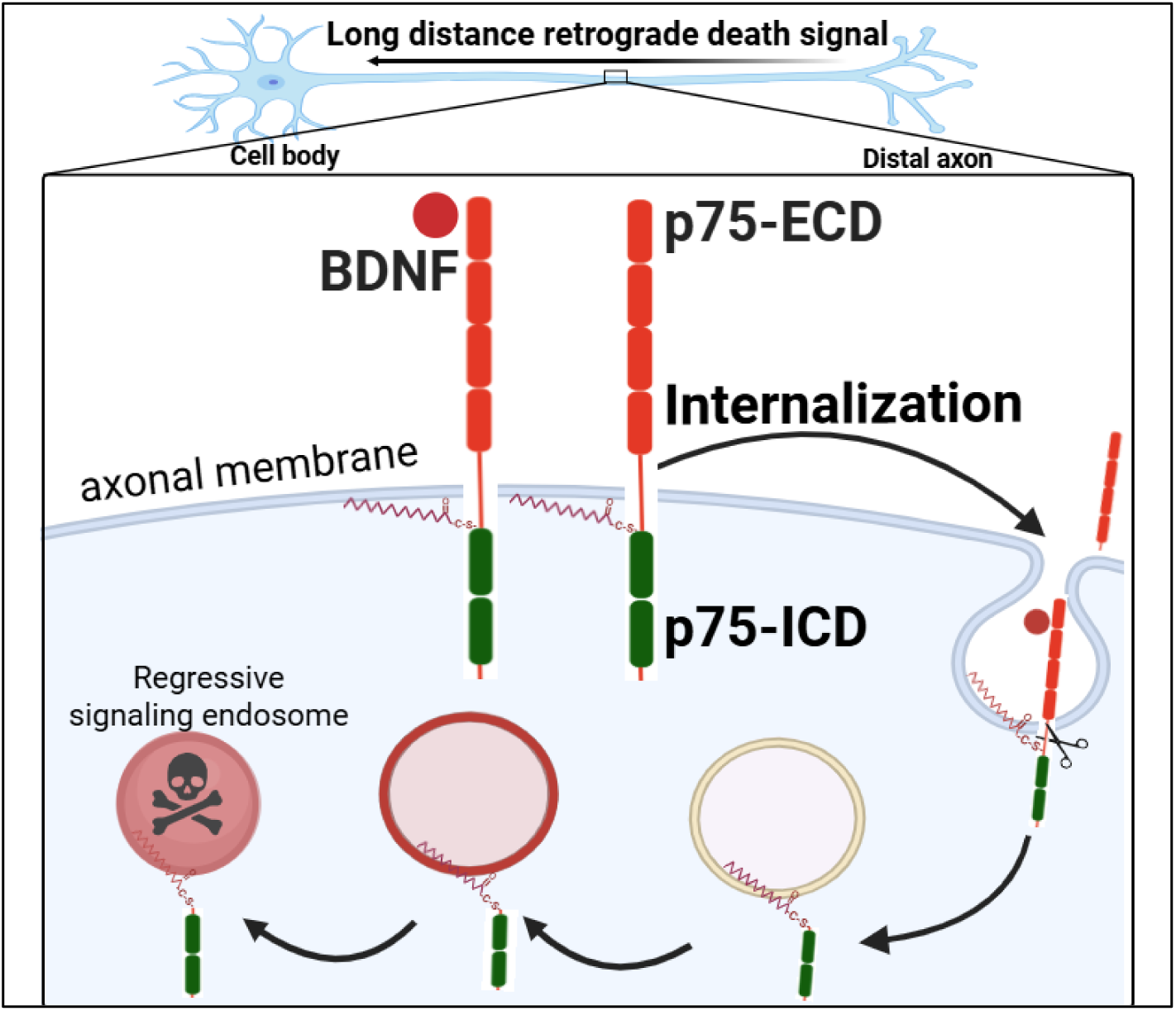

The intracellular domain (ICD) of neurotrophin receptor p75NTR is a long-distance death signal generated by its local cleavage in axons. Palmitoylation at conserved juxtamembrane Cysteine residue is necessary for ligand induced receptor internalization and provides a tether for attaching the cleaved p75ICD to regressive signaling endosomes leading to retrograde neuron death.

## Introduction

Assembly of the vertebrate nervous system depends on a tightly regulated balance between *progressive* events that stabilize neural connections and *regressive* events that remove supernumerary neurons, axons or synapses (Hamburger & Levi-Montalcini, 1949; Oppenheim, 1991). Target-derived neurotrophic factors such as the neurotrophins, including nerve growth factor (NGF), brain-derived neurotrophic factor (BDNF) and Neurotrophin-3 and -4 (NT3, NT4) signal through cognate receptor tyrosine kinases of the Trk family to promote survival, growth and synapse formation (Lewin & Barde, 1996). These pro-survival cues must be conveyed over long distances, often several centimeters in developing mammals, from distal axons back to the soma in the form of “signaling endosomes” (Harrington & Ginty, 2013; Grimes *et al*, 1996). Disruption of long-distance trophic signaling contributes to the pathogenesis of diverse neurodegenerative diseases including amyotrophic lateral sclerosis (ALS) and Alzheimer’s disease (Salehi *et al*, 2006; Perlson *et al*, 2009; Sleigh *et al*, 2019).

In parallel with these progressive pathways, developing neurons are exposed to long-distance *regressive* cues that instruct cell death, axon pruning or synapse restriction when innervation is inappropriate or trophic support is limiting (Deppmann *et al*, 2008; Pathak *et al*, 2021; Singh *et al*, 2008). The low-affinity tumour-necrosis-factor-receptor super-family member p75 neurotrophin receptor (p75NTR) is central to regressive signaling. In sympathetic neurons, binding of BDNF or the pro-neurotrophins, triggers regulated intramembrane proteolysis: ADAM17 first sheds the extracellular domain, then γ-secretase liberates the intracellular domain (ICD), which travels retrogradely in a dynein-dependent manner to activate apoptotic transcriptional programs in the soma (Kenchappa *et al*, 2006; Pathak *et al*, 2018; Escudero *et al*, 2014). Retrograde p75NTR signaling is required for developmental apoptosis of sympathetic and basal-forebrain cholinergic neurons and is re-engaged after injury or in neurodegenerative contexts (Dasgupta *et al*, 2024; Holland *et al*, 2016).

Relatively few regressive signaling molecules have been identified that retrogradely transport on axonal endosomes for long-distance communication. The best-characterized example is the MAP3K DLK, which must be *palmitoylated* to partition into retrograde endosomes, bind its scaffold JIP3 and activate JNK3 (Holland *et al*, 2016; Niu *et al*, 2022). In contrast, the mechanism by which p75NTR communicates over similar distances is largely unknown. Unlike Trk receptors, which remain membrane-anchored within signaling endosomes during retrograde transport, p75NTR cleavage releases a soluble ICD. This poses a biophysical puzzle: how does this soluble fragment remain associated with the endosomal surface to engage the dynein motor complex? One attractive solution is that palmitoylation of a conserved juxtamembrane cysteine transiently tethers p75NTR (or its ICD-containing C-terminal fragment) to endosomal membranes, thereby licensing engagement with dynein-driven retrograde carriers. Supporting this hypothesis, early biochemical studies detected p75NTR palmitoylation in PC12 cells, and inhibiting palmitoylation attenuated sensory-neuron death induced by p75NTR overexpression (Barker *et al*, 1994; Underwood *et al*, 2008). Whether, and how, palmitoylation regulates p75NTR endocytosis, endosomal sorting or long-distance degenerative signaling in primary neurons remains an open question.

Herein we asked whether palmitoylation of the conserved juxtamembrane cysteine (C281 in mouse p75NTR) could provide the transient membrane tether that enables endosomal sorting and long-distance transport of the otherwise soluble p75ICD. To test this idea, we: 1. Mapped p75NTR palmitoylation in primary sympathetic neurons and established its strict dependence on C281. 2. Visualized receptor trafficking in compartmentalized cultures to determine whether loss of palmitoylation blocks ligand-induced internalization and retrograde movement of p75ICD-positive endosomes. 3. Quantified functional consequences by measuring retrograde apoptosis *in vitro* and developmental neuron loss in a C281A *knock-in* mouse. Together, our findings identify palmitoylation as a critical mechanism that facilitates p75NTR inclusion in signaling endosomes and thereby enables long-distance regressive signaling.

## Results

### P75NTR is palmitoylated in neurons at a conserved cysteine residue in the intracellular domain

Previous biochemical analysis showed p75NTR palmitoylation in the neural cell line PC12 and transfected COS cells (Barker *et al*, 1994), however the cellular effects of p75NTR palmitoylation remained unclear. Interestingly the single cysteine residue to which the palmitate moiety is attached is highly conserved from fish to humans as shown in figure 1 A. The resin associated capture of acylated proteins and immunoblotting (Acyl RAC assay), showed robust palmitoylation of p75NTR in COS 7 cells transfected with the wildtype receptor. As demonstrated in figure 1C, the signal intensity for palmitoylated p75NTR decreased with a reduction in WT-p75NTR expression and increase in p75NTR^Δpalm^ mutant receptor where Cysteine279 was replaced with Alanine in rat p75NTR. The lack of palmitoylation in the p75NTR^Δpalm^ mutant confirmed that p75NTR is palmitoylated only at this particular cysteine residue. Sympathetic neuron death requires p75NTR cleavage and the release of the intracellular domain (ICD) fragment (Kenchappa et al., 2006), thus we asked whether palmitoylation can modify the cleaved intracellular domain fragment, hereafter referred to as p75ICD. We expressed either WT- or C279A mutant p75ICD fragment in COS7 cells and detected WT p75ICD palmitoylation, suggesting that the residues within this fragment are sufficient for palmitate addition (Figure 1D). We also probed p75NTR palmitoylation in lysates of sympathetic neurons cultured from superior cervical ganglion isolated from postnatal rats. As shown in figure 1E, endogenous p75NTR was palmitoylated in sympathetic neurons during early postnatal development.

**Figure 1.**
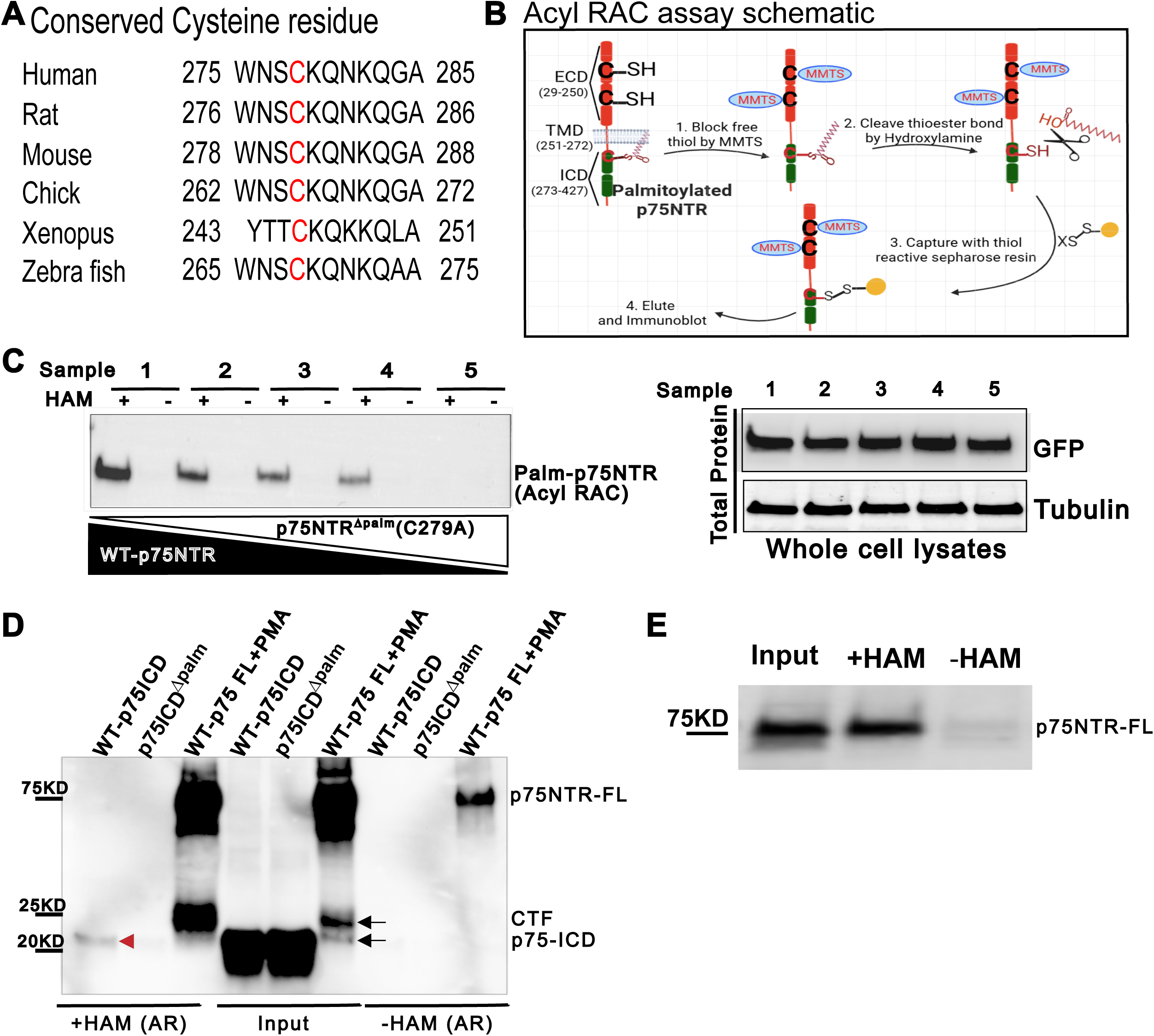
p75NTR palmitoylation at conserved Cysteine residue. (A) Comparison of protein sequences around the conserved Cysteine (C) residue from the indicated species. (B) Schematic for Acyl RAC assay used to determine p75NTR palmitoylation. (C) Representative images showing palmitoylated p75NTR detected by immunoblotting with GFP antibody after Acyl-RAC assay (Left) and total protein levels of respective samples numbered 1 through 5 (right). Whole cell lysates were prepared from COS7 cells transfected with either WT-p75NTR or palmitoylation resistant p75NTR^Δpalm^ mutant construct in decreasing levels of WT- (2ug to 0ug) and increasing levels of p75NTR^Δpalm^ (0 to 2 ug) construct as indicated in samples 1 through 5. After blocking the free thiols in the cysteine rich extracellular domain of p75NTR by methyl methane thiosulfonate (MMTS), hydroxylamine (HAM) was used to cleave the thioester bond, followed by capture of released thiol by Sepharose resin, elution and immunoblotting as indicated in the schematic in (B). Respective samples treated (+) or not treated (-) with HAM were loaded in adjacent lanes for clear comparison. (Right) Whole cell lysates immunoblotted with GFP and Tubulin showing similar levels of p75NTR-GFP expression. Compare lanes 1 and 5 in blots on the left and right. Note the complete absence of signal in cells expressing only the mutant p75NTR^Δpalm^. (D) Representative image showing palmitoylation of the intracellular domain (ICD) of p75NTR by immunoblotting with an antibody against p75ICD after Acyl-RAC assay. COS7 cells were transfected with WT-p75ICD only, palmitoylation resistant p75ICD only (p75ICD^Δpalm^) or full length WT-p75NTR-HA. COS7 cells expressing full length p75NTR-HA were treated with 1µM PMA for 4 hours. Prior to PMA treatment, COS7 cells were pretreated with 5µM proteasome inhibitor Clasto-lactacystin β-lactone (CL). Note the band for WT palmitoylated p75ICD in lane 1 compared to mutant p75ICD^Δpalm^ in lane 2. Input indicates 20 ug of total protein after treating the lysates with MMTS. (E) Immunoblot after Acyl RAC assay showing palmitoylation of endogenous p75NTR present in sympathetic neurons cultured from WT rat superior cervical ganglia.

### Palmitoylation is required for p75NTR internalization and regressive signaling endosome biogenesis in axons

We previously demonstrated that treatment of distal axons with BDNF induced local cleavage of p75NTR, resulting in retrograde transport of the p75ICD, ultimately leading to sympathetic neuron death (Pathak *et al*, 2018). Thus, we hypothesized that palmitoylation facilitates this trafficking of p75ICD towards the neuronal soma. To interrogate the requirement of this lipid modification for axonal transport of p75ICD, we cultured wildtype primary sympathetic neurons in microfluidic chambers, which allows separation of the cell bodies and proximal axons from the distal axons (Fig. 2A). After culturing the neurons in NGF and allowing them to extend their axons into the distal chamber, we replaced the NGF with potassium chloride (KCl). Mild depolarization keeps the neurons alive but avoids the complexity of exposure to a neurotrophin (NGF) that binds both TrkA and p75NTR. The distal axons were then either treated with BDNF, which only binds to the p75NTR in these neurons, or BDNF along with palmitoylation inhibitor, 2-bromopalmitate (2Br), which blocks acylation. The soma compartment was left untreated. Control cultures were maintained in KCl both at the axon and soma compartments. After 2 hours, the cultures were fixed and immunostained using an antibody against the intracellular domain of p75NTR. This antibody cannot distinguish between the cleaved p75ICD and the full-length p75NTR; however, we expect a majority of puncta to be p75ICD as BDNF treatment of distal axons leads to the cleavage of axonal p75NTR as reported earlier (Pathak et al, 2018). The distal axons were imaged with Airyscan mode on Zeiss confocal microscope for clear visualization and quantification of the p75ICD puncta. As demonstrated in figure 2B and 2C, p75ICD puncta were significantly increased in distal axons following BDNF treatment, likely reflecting trafficking endosomes. Importantly, the formation of these puncta was completely blocked by 2Br applied to the distal axons (Fig. 2B & C). These results suggest that inhibiting p75NTR palmitoylation only in the distal axons prevented the BDNF-induced vesicular localization of p75ICD, however we cannot completely exclude the presence of endogenous full-length receptors in these axonal vesicles.

**Figure 2.**
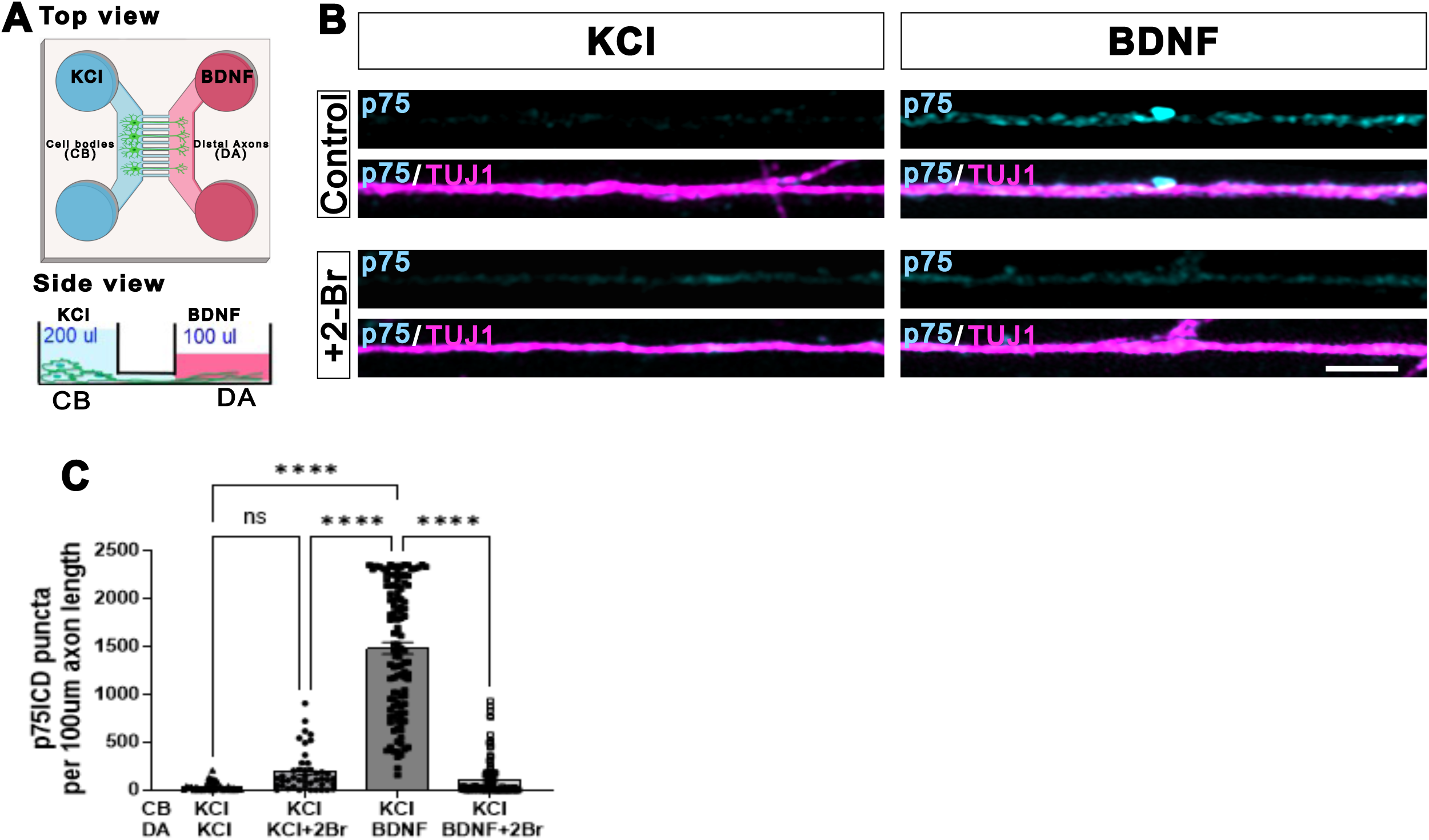
Palmitoylation is required for BDNF-induced p75ICD endosome biogenesis in distal axons. (A) Schematic of microfluidic devices (MFDs) fluidically isolating cell bodies (CB) from distal axons (DA) by differential media volume in the left (blue, 200µl) and right (pink, 100µl) chambers. Sympathetic neurons from WT rats were cultured in media with 20ng/ml NGF, after the axons had reached the DA chamber, NGF was removed and neurons were maintained in 12.5 mM KCl. DA were treated with BDNF in presence or absence of palmitoylation inhibitor 2-bromo palmitate (2Br) for 2 hours. Cell bodies were kept in KCl. Fixed neurons were immunostained with an antibody against p75ICD (cyan) and neuronal marker TUJ1 (magenta). To quantify the puncta only the axons reaching the DA chamber were imaged on Zeiss LSM980 confocal microscope on Airyscan mode. Immunostaining for p75ICD (cyan) revealed an increase in vesicular puncta after BDNF treatment, which was blocked by 2Br treatment. Scale bar: 2 µm. (B) Quantification of p75ICD puncta per 100 µm axon length. Each data point indicates an individual axon. Inhibition of palmitoylation by 2Br treatment of DA prevents the BDNF-induced increase in axonal puncta. Data represent mean ± SEM from more than 100 axons from two different experiments. The data was analyzed by 2-way ANOVA using Tukey’s multiple comparison test. ****P<0.0001, ns: non significant.

Pharmacological inhibitors like 2Br broadly inhibit palmitoylation of all proteins, thus, to assess the effects specifically due to palmitoylation of p75NTR we expressed either WT or palmitoylation resistant p75NTR^Δpalm^ with GFP fused to the C-terminus of p75NTR in sympathetic neurons using lentivirus mediated transduction. Both the WT and p75NTR^Δpalm^ receptors were expressed and trafficked to the plasma membrane to a similar extent based on a surface biotinylation assay (Fig. 3A). We also detected both forms of the receptor in the axons by immunostaining (Fig. 3B). These results indicate that blocking p75NTR palmitoylation does not affect anterograde trafficking to the axon or the cell surface. Neurons expressing the WT or mutant receptor were then grown in microfluidic chambers as above and distal axons treated with BDNF or left untreated. After 2 hours, the neurons were fixed and the amount of GFP puncta in the axons was quantified. Wild type p75NTR accumulated in axonal vesicles following BDNF treatment. However, the level of vesicular p75NTR^Δpalm^ after BDNF treatment was significantly lower (Fig. 3B & C), supporting the idea that p75NTR palmitoylation is required for axonal endosome formation in response to BDNF.

**Figure 3.**
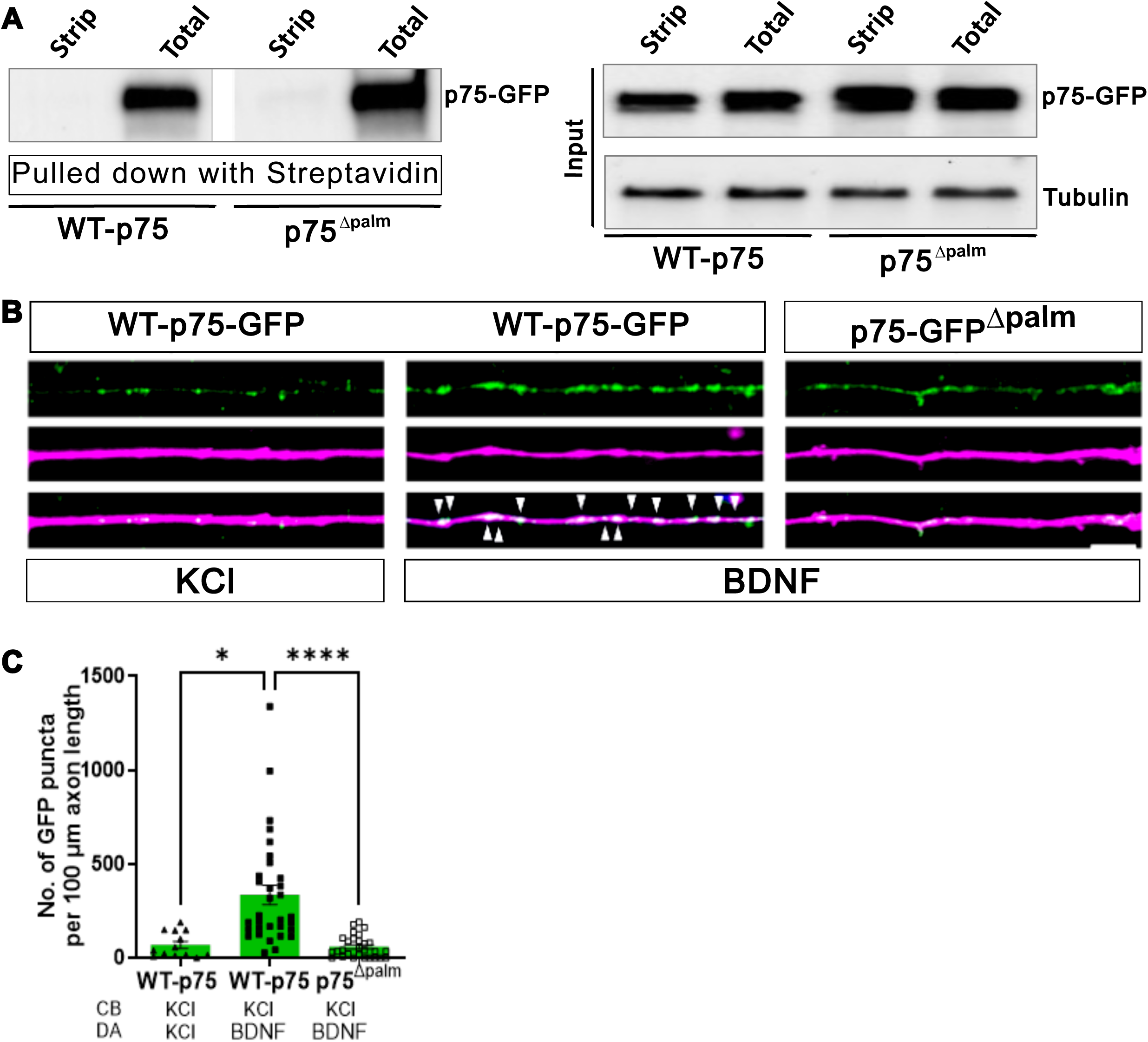
Palmitoylation does not affect p75NTR surface expression but it is required for BDNF-induced vesicular localization of p75NTR. (A) Primary neurons were grown in mass cultures and were infected with lentivirus to express either WT-p75NTR-GFP or mutant p75NTR^Δpalm^-GFP. Following biotinylation of surface proteins, immunoblotting of streptavidin precipitated samples and input lysates was done with anti-GFP antibody, which showed equal surface expression of WT-p75NTR and p75NTR^Δpalm^ (left). Input samples were immunoblotted with antibodies against GFP and Tubulin as controls (right). (B) Representative images of axons of primary sympathetic neurons cultured from p75^-/-^ animals in MFDs and infected with lentivirus to express WT-p75NTR or p75NTR^Δpalm^-GFP. DA were treated with BDNF for 2 hours as indicated in (B) and (C). To quantify the axonal puncta only the axons reaching the DA chamber were imaged on Zeiss LSM980 confocal microscope on Airyscan mode. White arrowheads indicate BDNF induced p75NTR accumulation in axonal vesicles in neurons expressing WT-p75NTR GFP, which was significantly lower in neurons expressing p75NTR^Δpalm^ mutant. (C) Quantification of GFP-positive axonal puncta. Each data point indicates an individual axon, analyzed from two independent experiments. Data represent mean ± SEM which was analyzed by 2-way ANOVA using Tukey’s multiple comparison test. ****P<0.0001, *P=0.0239.

As a first step in endosome formation, the receptor must be internalized; therefore, we asked whether palmitoylation affects ligand-induced p75NTR internalization. We expressed WT p75NTR or p75NTR^Δpalm^ with the GFP tag in sympathetic neurons and evaluated endocytosis using a pHrodo-labeled antibody to the extracellular domain of the receptor. To avoid any influence of the endogenous p75NTR, we used neurons from p75NTR^-/-^ mice and expressed WT or p75NTR^Δpalm^-GFP using lentivirus. The neurons expressing WT or p75NTR^Δpalm^, grown in mass cultures, were treated with or without BDNF for at least 2 hours together with a pHrodo-labeled antibody that does not interfere with ligand binding. The neurons were then imaged live and receptor internalization in pHrodo labeled acidic vesicles was quantified by scoring GFP+ and pHRodo+ signal, both in the cell body and in the axons as shown in figure 4B and 4C. Although BDNF stimulated significant internalization of WT p75NTR, there was no endocytosis detected of the p75NTR^Δpalm^ mutant. These results suggest that lipid modification of p75NTR is required for ligand-mediated internalization. Furthermore, to assess p75NTR internalization specifically in axons, we cultured the p75NTR^-/-^ neurons in microfluidic chambers and expressed either WT or p75NTR^Δpalm^ by lentivirus transduction and quantified BDNF induced axonal puncta for WT and mutant p75NTR-GFP. As demonstrated in figure 4D and 4E, BDNF treatment exclusively on the axons also showed reduced pHrodo-labeled p75NTR puncta for the palmitoylation resistant mutant compared to WT-p75NTR.

**Figure 4.**
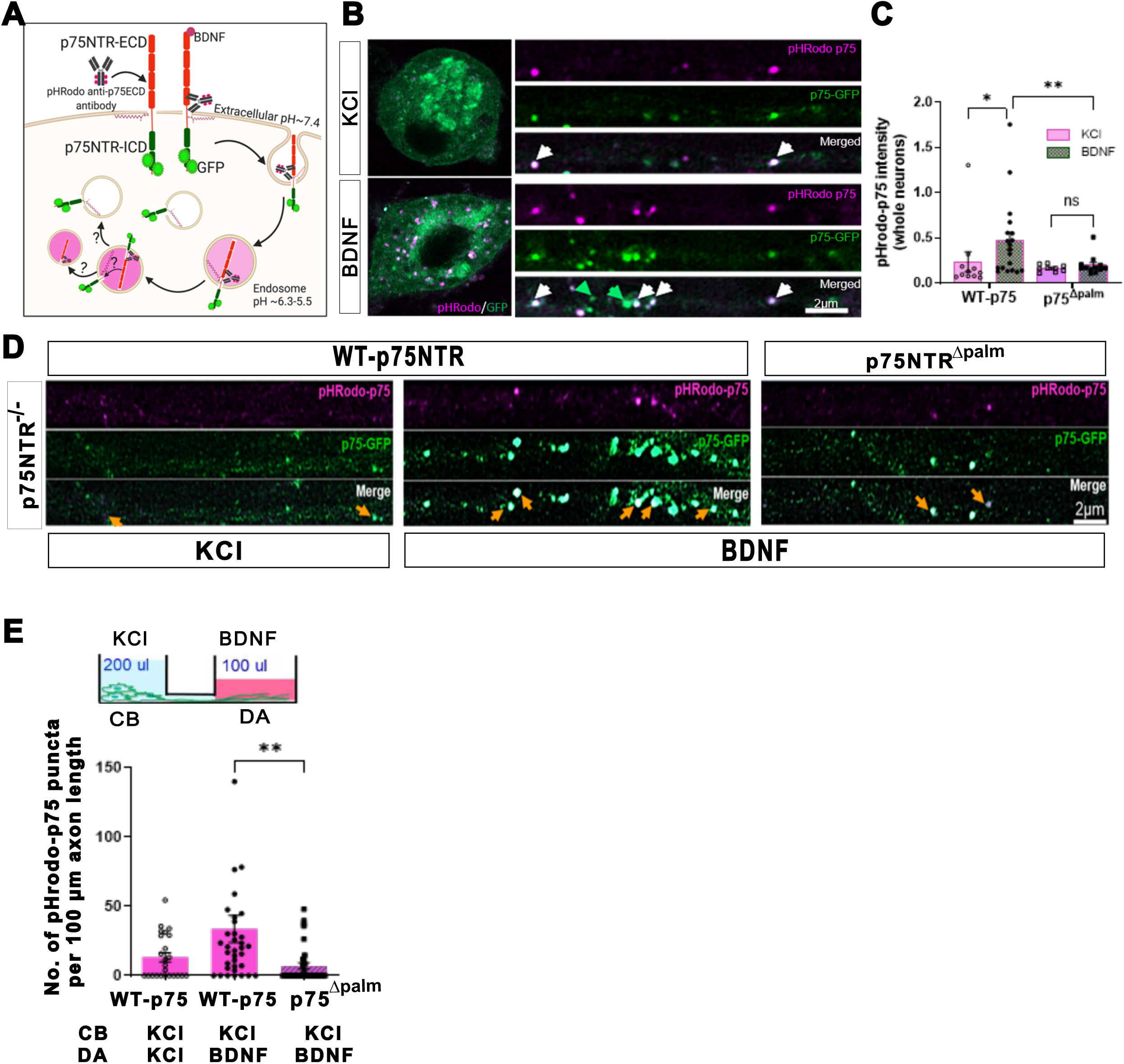
Palmitoylation is required for ligand-induced p75NTR internalization. (A) Schematic of the pHrodo-p75NTR internalization assay using the antibody against the extracellular domain of p75NTR conjugated with pH sensitive dye pHrodo which fluoresces red only at acidic pH of endosomes. (B) Representative images of P75^-/-^ neurons in mass culture expressing WT-p75NTR-GFP by lentivirus transduction and labeled with pHrodo-p75NTR antibody and treatment of whole neurons with BDNF for at least two hours to induce internalization or left untreated in KCl as control. Neurons were imaged live on Zeiss LSM980 confocal microscope on Airyscan mode. White arrows indicate pHrodo+/GFP+ double labeled axonal puncta while green arrows indicate GFP only puncta. Cell bodies and axons are shown separately for clear visualization of pHrodo and GFP puncta. (C) Quantification of internalized p75NTR in pHrodo-positive vesicles in whole neurons expressing either WT or p75NTR^Δpalm^ mutant. Upon BDNF treatment WT-p75NTR showed robust internalization measured as an increase in pHrodo intensity. WT p75NTR undergoes robust internalization with BDNF, whereas the palmitoylation resistant p75NTR^Δpalm^ mutant fails to internalize in pHrodo labeled acidic vesicles. Data represent mean ± SEM from three independent experiments which was analyzed by 2-way ANOVA using Tukey’s multiple comparison test. **P<0.01, *P=0.015. (D) p75NTR^-/-^ neurons were cultured in MFDs and expressed either WT- or p75NTR^Δpalm^-GFP by lentivirus transduction to quantify p75NTR internalization exclusively in distal axons following BDNF treatment. Yellow arrows indicate pHrodo+GFP+ double labeled puncta in axons. (E) Quantification of pHrodo-positive axonal puncta. Each data point indicates an individual axon. Data represent mean ± SEM from two independent experiments which was analyzed by 2-way ANOVA using Tukey’s multiple comparison test. **P<0.01.

### Palmitoylation is not required for the regulated proteolysis of p75NTR and does not alter the stability of its cleavage fragments

It has been suggested that p75NTR cleavage by γ-secretase occurs in endosomes, following receptor internalization (Urra *et al*, 2007). Given our results indicating that p75NTR palmitoylation is required for endocytosis, led us to question whether receptor cleavage would be affected. Therefore, we expressed either WT or palmitoylation resistant mutant receptor in COS cells, which were then treated with phorbol methyl ester (PMA) to induce cleavage. A proteasome inhibitor, clastolactacystin, was included because the p75ICD is typically quickly degraded (Kanning *et al*, 2003; Jung *et al*, 2003). We found both WT and p75NTR^Δpalm^ underwent proteolytic cleavage to a similar extent, with more p75ICD generated from the p75NTR^Δpalm^ mutant initially (Fig. 5A-E) which later plateaued. Therefore, palmitoylation is not required for the regulated proteolysis of p75NTR.

**Figure 5.**
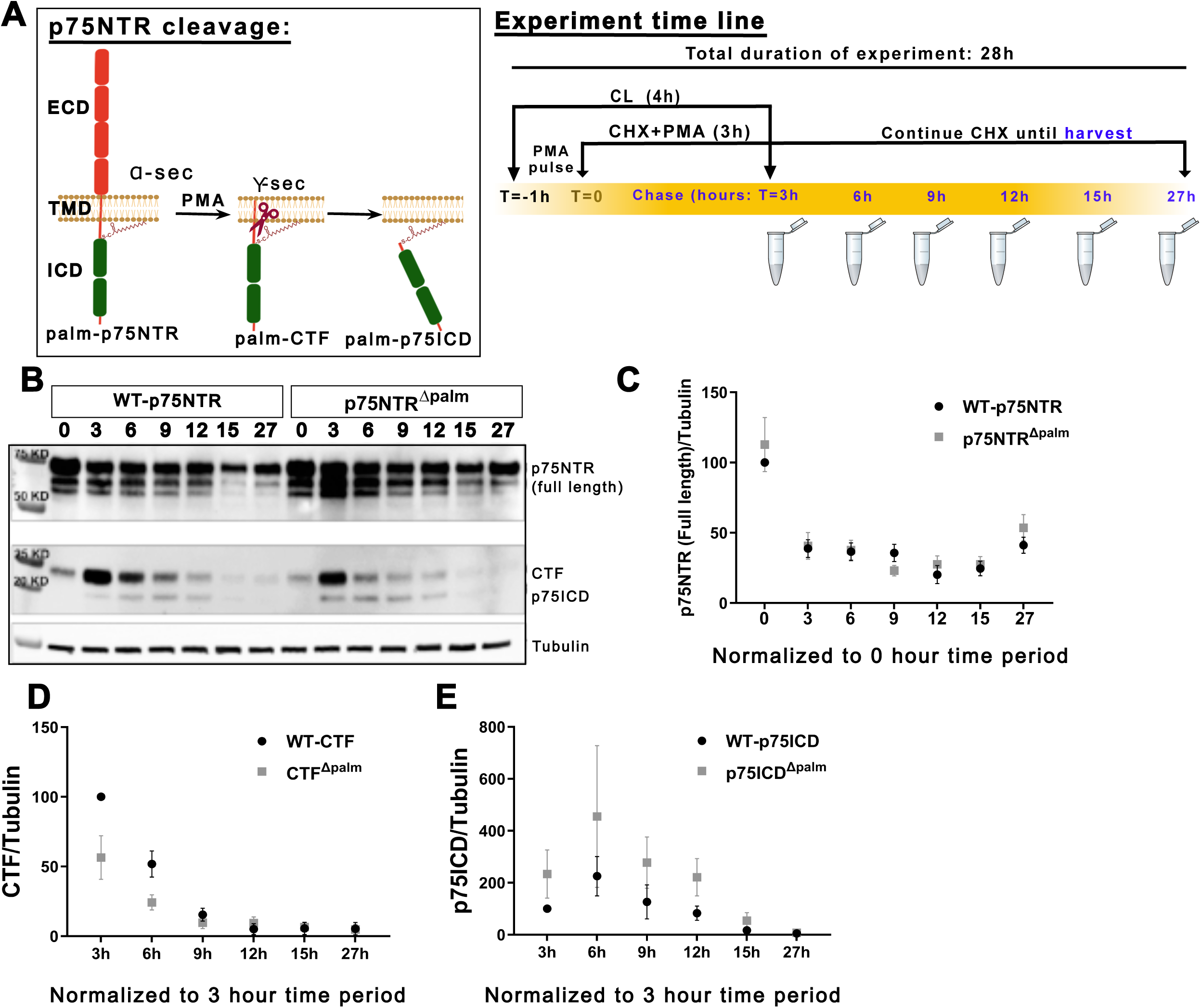
Palmitoylation does not affect p75NTR stability, nor it is required for its regulated proteolysis. (A) (Left) Schematic showing regulated intramembrane proteolysis of p75NTR by α- and γ-secretase to release the intracellular domain, p75ICD. (Right) Experiment timeline to investigate p75NTR cleavage in WT-p75NTR or p75NTR^Δpalm^ expressing COS7 cells. 1µm PMA (pulse) induced the cleavage followed by biochemical chase after several hours of cycloheximide (CHX) treatment as indicated in blue. 5µg/ml CHX treatment blocked new synthesis of p75NTR. Proteasome inhibitor Clasto-lactacystin β-lactone (CL) treatment at 5µM inhibited degradation of C-terminal fragment (CTF) and p75ICD. (B) COS7 cells were transfected with WT-p75NTR or p75NTR^Δpalm^ and were treated as indicated in the pulse chase schematic shown in (A). Whole cell lysates were immunoblotted with an antibody against the intracellular domain of p75NTR. Cropped images of different exposures of the same blot are shown for clear visibility of bands (top and middle). Immunoblotting with Tubulin confirmed equal loading. (C-E) Graphs showing p75NTR stability and cleavage after chase duration. Full length p75NTR was normalized to WT-p75NTR at time point zero, whereas CTF and p75ICD was normalized to wildtype CTF and p75ICD at three hour time period as these cleavage fragments were detectable only after three hours of PMA treatment, not at time point zero. Shown here are the mean values from four different experiments. Statistical analysis was done by 2-way ANOVA with Sidak’s multiple comparison test.

The dramatic decrease in detectable internalized p75NTR^Δpalm^ relative to WT p75NTR could also be the result of altered stability of the receptor or its cleavage products. Palmitoylation can regulate protein stability; for example, flotillin (McClellan *et al*, 2024) and SynDIG1 (Kaur *et al*, 2016), which modulates synaptic AMPA receptor content, are stabilized by lipidation. Therefore, to determine whether the stability of full length p75NTR or its proteolytic products are affected by palmitoylation, we examined the processing and stability of the WT receptor compared to the p75NTR^Δpalm^ mutant. COS cells were transfected with WT p75NTR or p75NTR^Δpalm^ and after 48 hours of transfection, the cells were treated with the proteasome inhibitor clastolactacystin (CL) to prevent degradation of the intracellular domain fragments, followed by phorbol methyl ester treatment to induce receptor cleavage and cycloheximide to block new protein synthesis. After 3 hours, the CL and PMA were removed, but cycloheximide treatment was continued. We reasoned that phorbol ester would induce cleavage of the receptor and the fragments would accumulate without any degradation due to proteasome inhibition. Upon removal of PMA and CL, the cleavage of p75NTR would cease, and the cleaved receptor fragments could be degraded. No new receptor would be synthesized in the presence of cycloheximide. As shown in figure 5B-E, we did not observe any notable difference between WT p75NTR and p75NTR^Δpalm^ upon comparison of the stability of the full length receptor or its cleavage products, C-terminal fragment (CTF) and p75ICD. Thus, palmitoylation of p75NTR does not appear to affect the stability of the receptor or its cleavage products.

### P75NTR palmitoylation is required for long distance regressive signaling in cultured sympathetic neurons

Given that p75NTR internalization and endosome formation were prevented by blocking receptor palmitoylation, we hypothesized that p75NTR mediated neuron apoptosis following retrograde signaling in axons would also be blocked. To test this hypothesis, wildtype rat sympathetic neurons were cultured in microfluidic chambers and the distal axons treated with or without 2Br to block local palmitoylation along with BDNF or left untreated. After 48 hours, apoptosis of the cell bodies was quantified. The addition of 2Br in the distal axons prevented BDNF induced cell death (Fig. 6A).

**Figure 6.**
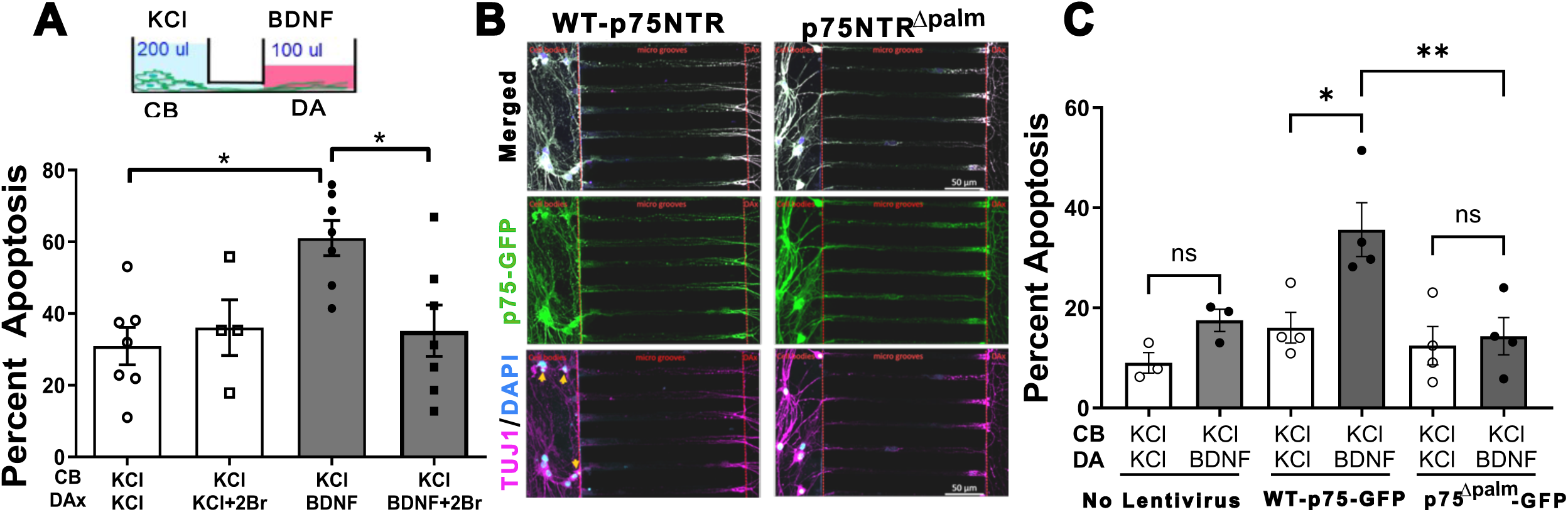
P75NTR palmitoylation is necessary for long distance degenerative signaling in neurons. (A) Wildtype primary neurons were cultured in MFDs in media with 20ng/ml NGF, after the axons reached the DA chamber, NGF was removed and neurons were maintained in 12.5 mM KCl. DA were treated with BDNF in presence or absence of palmitoylation inhibitor 2-bromo palmitate (2Br) for 48 hrs and CB were kept in KCl as indicated. Fixed neurons were stained with TUJ1 and DAPI to score pyknotic nuclei as a measure of apoptosis. Depicted are the means ± SEM for n = 4; **, p < 0.01; 2-way ANOVA. (B) Representative images of p75NTR^-/-^ neurons cultured in MFDs and infected with lentivirus to express WT or palmitoylation resistant p75NTR^Δpalm^. After the expression of p75NTR, NGF was removed and DA were treated with BDNF and scored for apoptosis as indicated and mentioned above. Yellow arrows indicate DAPI stained pyknotic nuclei in the CB chamber. (C) Expression of p75NTR^Δpalm^ (right) prevents BDNF induced apoptosis as seen in p75NTR^-/-^ neurons (left). Depicted are the means ± SEM for n >3; **, p < 0.01; 2-way ANOVA with Tukey’s multiple comparison test.

To more specifically assess the influence of p75NTR palmitoylation and avoid the complication of the endogenous receptor, we expressed WT p75NTR or p75NTR^Δpalm^ in sympathetic neurons isolated from p75NTR^-/-^ mice and cultured them in the microfluidic chambers. The distal axons were treated with BDNF for 48 hours or left untreated in KCl as controls and neuronal apoptosis was scored. As expected, expression of WT p75NTR facilitated BDNF-mediated retrograde apoptotic signaling. However, neuronal death was completely blocked in the neurons expressing p75NTR^Δpalm^ (Fig. 6B & C). These results suggest that blocking palmitoylation of p75NTR prevents retrograde BDNF induced retrograde death signaling in sympathetic neurons.

### p75NTR palmitoylation is required for receptor internalization and neuron death *in vivo*

To evaluate the effects of p75NTR palmitoylation *in vivo*, we generated mice carrying the C281A mutation (the mouse equivalent of rat C279A) using CRISPR CAS9. This DNA modification introduced a novel PvuII restriction site that provided a method for easy genotyping (Fig. 7A). We backcrossed these mice for 7 generations, to avoid any off-target effects and also confirmed the genotypes by Sanger sequencing. By immunoblotting the lysates from superior cervical ganglion (SCG), we confirmed that the mutant receptor was expressed to a similar extent as WT p75NTR (Fig. 7B) and as shown in figure 7C, the Acyl RAC assay verified that p75NTR was not palmitoylated in the C281A (p75NTR^Δpalm/Δpalm^) mutant mice.

**Figure 7.**
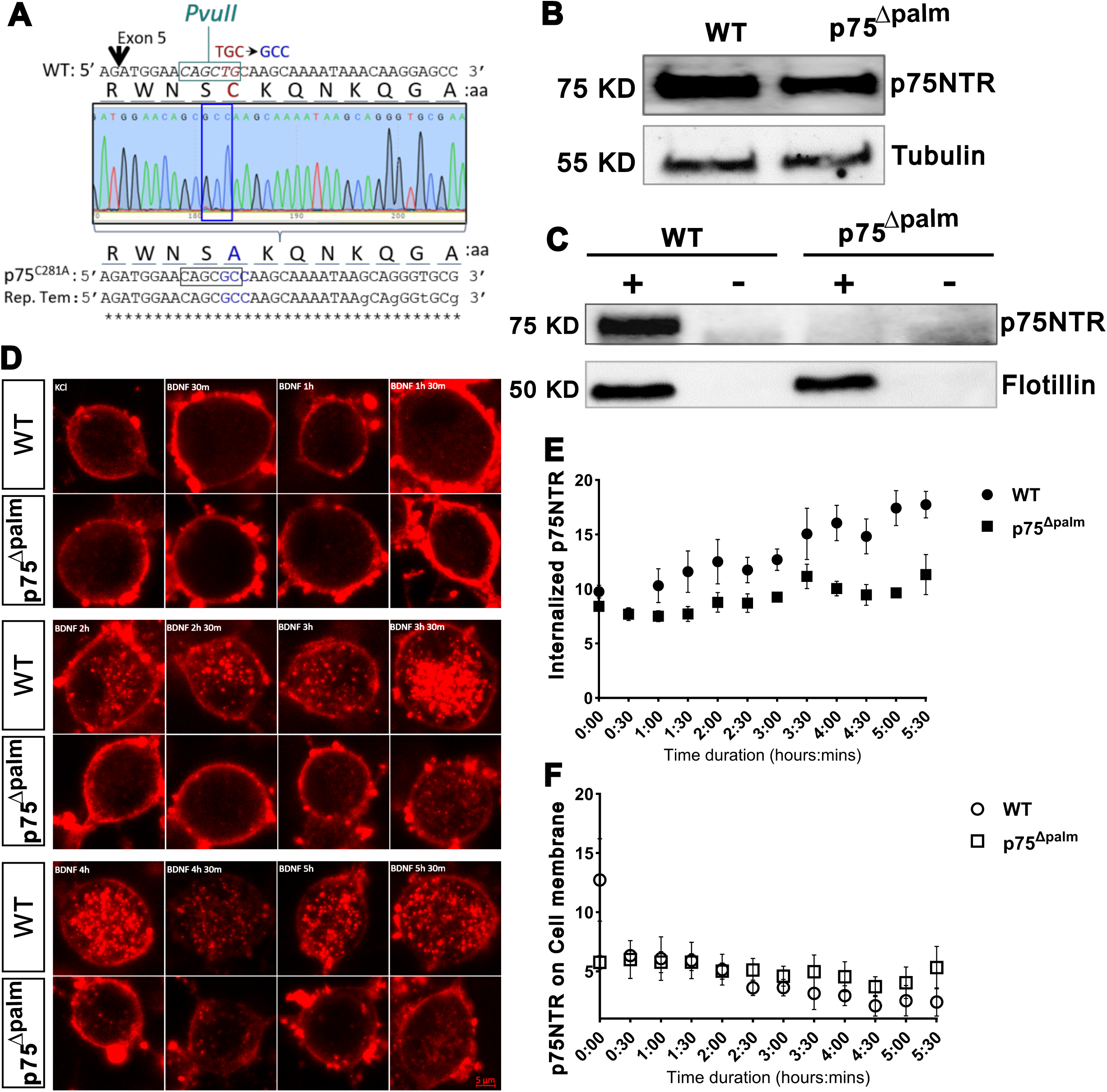
Palmitoylation is required for endogenous p75NTR internalization in primary sympathetic neurons. (A) Amino acid and DNA sequences of the region around Cysteine 281 and the chromatogram showing strategy for p75NTR^Δpalm/Δpalm^ mice generation by CRISPR/Cas9. TGC (Cys) to GCC (Ala) mutation was confirmed by PvuII digestion followed by Sanger sequencing of the amplicons including the mutant sequence. (B) Immunoblot of the whole cell lysates prepared from the sympathetic ganglia of WT and p75NTR^Δpalm/Δpalm^ mice showed similar p75NTR expression. (C) Acyl RAC assay demonstrated complete absence of p75NTR palmitoylation in p75NTR^Δpalm/Δpalm^ mice. Samples were also immunoblotted for flotillin as a control for palmitoylation. (D) Representative images showing BDNF induced p75NTR immuno-endocytosis in live sympathetic neurons cultured from WT- or p75NTR^Δpalm/Δpalm^ mice. Sympathetic neurons were incubated with an antibody against p75NTR extracellular domain conjugated with ATTO550 at 4°C and then treated with 200 ng/mL BDNF at 37°C for at least 30 minutes to several hours as indicated. The labelled neurons were imaged live and Z-stack images were collected. Images were taken at a 30 minute interval until 5 hours and 30 minutes. Shown here are collapsed Z-stack images with equal number of slices in each neuron. The duration of BDNF treatment is indicated on the top left corner of the image panel for WT-neurons and images for p75NTR^Δpalm/Δpalm^ neurons are shown underneath the WT neuron at corresponding time point. (E & F) Quantification of internalized and non-internalized p75NTR which was present on the neuronal membrane following BDNF treatment. The internalized p75NTR corresponds to the intracellular fluorescence in Z-stacks with equal number of z-slices. Shown is the average internalized fluorescence ± SEM for n>5 experiments. A 2-way Anova with Tukey’s multiple comparison test showed significant difference in p75NTR internalization between WT and p75NTR^Δpalm/Δpalm^ neurons (E). Over the time course, there was a significant increase in BDNF induced WT-p75NTR internalization (p=0.0092), whereas there was no significant difference in p75NTR internalization in p75NTR^Δpalm/Δpalm^ neurons following BDNF treatment.

To determine whether p75NTR internalization was inhibited in the p75NTR^Δpalm/Δpalm^ mice, similar to what we observed when the mutant receptor was expressed in p75NTR^-/-^ neurons (Fig. 4), we isolated the sympathetic neurons from WT and p75NTR^Δpalm/Δpalm^ mice SCGs. The neurons were cultured for 24 hrs, then treated with a fluorescently labeled antibody to the extracellular domain of p75NTR. This antibody does not prevent ligand binding, thus BDNF binding to p75NTR was not affected. The neurons were imaged live for over 5 hours and internalized as well as cell surface p75NTR were quantified as shown in Fig. 7D-E. WT p75NTR internalized over a period of several hours with less remaining on the cell surface, similar to what has previously been reported (Bronfman *et al*, 2003). In contrast, the p75NTR^Δpalm/Δpalm^ neurons exhibited minimal receptor endocytosis and continued expression on the cell surface.

During development, p75NTR plays an essential role in orchestrating the balance between death and survival in sympathetic neurons (Majdan *et al*, 1997; Deppmann *et al*, 2008; Bamji *et al*, 1998). Deletion of the receptor results in less apoptosis and an excess of these neurons (Bamji *et al*, 1998; Kristiansen & Ham, 2014). Therefore, we investigated whether cell death was affected in the sympathetic neurons of the superior cervical ganglia of the p75NTR^Δpalm/Δpalm^ mice. Co-staining for cleaved caspase 3 and the neuronal marker, TuJ1, in the mice pups at postnatal day zero (p0) revealed a marked decrease in the degree of neuronal apoptosis, similar to the complete deletion of the receptor (Fig. 8A & B). These results indicate that lipid modification of p75NTR is required for apoptotic signaling *in vivo*.

**Figure 8.**
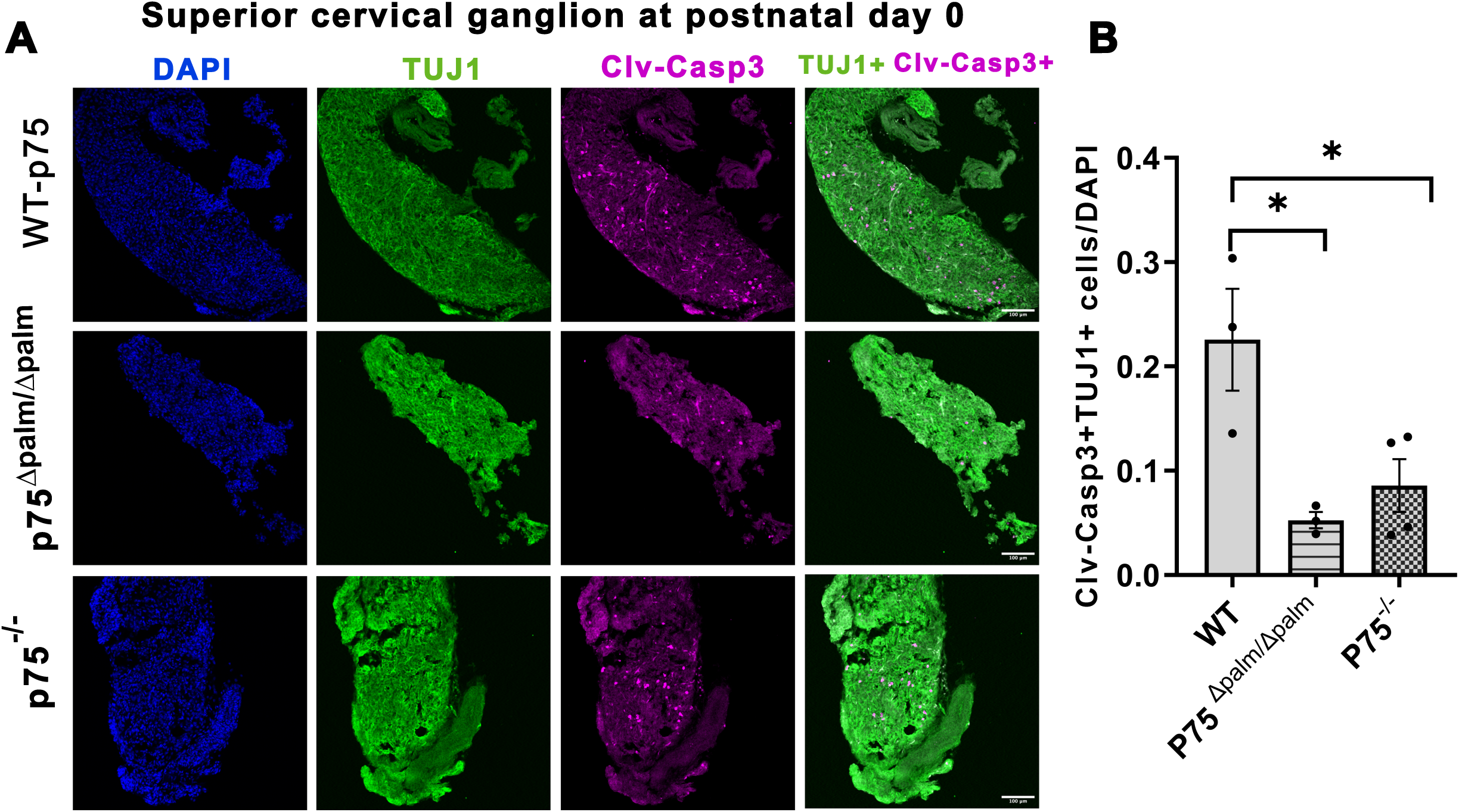
Sympathetic neuron death during development requires p75NTR palmitoylation. (A) Representative images showing neuron apoptosis in superior cervical ganglia of WT, p75NTR^-/-^ and p75NTR^Δpalm^ pups at postnatal day 0. Co-staining for cleaved caspase 3 and TuJ1 revealed significant decrease in apoptosis, similar to the p75NTR knockout mice. (B) Quantification of the cleaved caspase 3 and TuJ1 double positive neurons which were normalized to the total number of cells by DAPI staining. Depicted are the means ± SEM for n >3; *, p < 0.05; one-way ANOVA with Dunnett’s multiple comparison test.

## Discussion

The mechanism by which p75NTR regressive signaling molecules traverse long distances from axon terminals to the soma has remained an open question in developmental neurobiology. Here we demonstrate that palmitoylation of a single conserved cysteine residue (C281) in p75NTR is required for efficient ligand-induced internalization and long-distance retrograde apoptotic signaling. Using compartmentalized cultures, live-cell imaging, and a palmitoylation-deficient *knock-in* mouse, we show that loss of this lipid modification blocks ligand-induced internalization, strongly reduces formation of p75ICD-containing endosomes in distal axons, impairs retrograde signaling, and largely phenocopies p75NTR deletion with respect to developmental sympathetic neuron death. These findings address a longstanding biophysical challenge: how can a soluble intracellular domain fragment, liberated by regulated proteolysis, be conveyed over centimeter-scale distances? Our data support a model in which palmitoylation provides a transient membrane tether that anchors p75ICD to endosomal membranes, thereby enabling engagement with the retrograde transport machinery previously shown to require dynein (Pathak *et al*, 2018; Escudero *et al*, 2019). We identify C281 as the critical palmitoylation site in neurons, demonstrate that palmitoylation is required for ligand-induced internalization and retrograde death signaling, and provide genetic evidence from the C281A *knock-in* mouse that this modification is required for developmental apoptosis *in vivo*.

### Palmitoylation as a Molecular Tether for Long-Distance Regressive Signaling

The p75NTR receptor poses a unique challenge for long-distance signaling. Unlike receptor tyrosine kinases such as TrkA, which remain membrane-anchored throughout their retrograde journey (Harrington & Ginty, 2013; Grimes *et al*, 1996), p75NTR undergoes sequential proteolytic processing that liberates a soluble approximately 20 kDa intracellular domain (Kanning *et al*, 2003) (Jung *et al*, 2003; Kenchappa *et al*, 2006). Our discovery that this ICD can be palmitoylated when expressed in COS7 cells, building on early evidence of p75NTR palmitoylation in PC12 and COS cells (Barker *et al*, 1994), provides a potential mechanistic solution. The 16-carbon palmitate moiety could serve as a lipid anchor that maintains association with endosomal membranes despite loss of the transmembrane domain. We observe that palmitoylation persists on both the C-terminal fragment and ICD after cleavage in overexpression systems.

Alternative mechanisms could contribute to our findings. For example, palmitoylation might influence receptor clustering at the plasma membrane, adaptor protein binding, or endosomal sorting events. Thus, while our data support palmitoylation being required for receptor internalization as well as acting as a transient membrane anchor, we cannot yet distinguish whether this requirement arises at the level of receptor clustering, protein interaction, endosomal sorting, or all three.

### Context-Dependent Effects of Death Receptor Palmitoylation

In contrast to our findings, recent work by Ma et al (2026) found that loss of p75NTR palmitoylation in hippocampal neurons accelerated internalization via enhanced clathrin-mediated endocytosis and promoted lysosomal degradation, effects that supported a neuroprotective role in the 5xFAD Alzheimer’s disease model by enhancing APP degradation (Ma *et al*, 2026). Despite following a similar experimental paradigm using the antibody feeding internalization assay for p75NTR, the apparent discrepancy likely reflects context-dependence: different neuron types (sympathetic vs. hippocampal), different ligand contexts (BDNF-driven developmental death vs. chronic Aβ/APP exposure), and likely different receptor complexes (p75NTR alone or with Sortilin in sympathetic neurons vs. APP/p75NTR in hippocampal neurons). Together, these observations suggest that palmitoylation may license p75NTR entry into signaling competent carriers that support productive retrograde transport in sympathetic neurons (Underwood *et al*, 2008; Ma *et al*, 2026) while in other contexts serving as a retention signal that slows progression through the endolysosomal system.

Context-dependent effects extend to other death receptors. TNF receptor 1 (TNF-R1) is constitutively palmitoylated at C248, and ligand-induced depalmitoylation by APT2 facilitates internalization and the switch from pro-survival to pro-death signaling (Zingler *et al*, 2019). Similarly, palmitoylation enhances death signaling for Fas and TRAIL receptor DR4 (Chakrabandhu *et al*, 2007; Rossin *et al*, 2009). Importantly, the directionality differs: depalmitoylation promotes death signaling for TNF-R1, whereas palmitoylation is required for p75NTR mediated neuron death signaling. This underscores that S-acylation configures receptor-specific trafficking programs rather than acting as a universal “death switch” (Rabizadeh *et al*, 1993; Zingler *et al*, 2019). The dynamic and reversible nature of palmitoylation, coupled with the large family of DHHC enzymes and depalmitoylases with distinct expression patterns, provides a flexible system for spatially and temporally controlling death receptor function.

### Comparison to DLK: A Common Requirement for Lipid Modification in Regressive Signaling?

The requirement for palmitoylation of p75NTR parallels recent findings for another key regressive signaling molecule, the MAP3K dual leucine zipper kinase (DLK). Thomas and colleagues (2016) demonstrated that DLK palmitoylation at N-terminal cysteine is essential for its localization to vesicular compartments, interaction with the scaffold JIP3, and activation of JNK-dependent axon degeneration (Holland *et al*, 2016). More recently, the palmitoyl acyltransferase ZDHHC17 was identified as a key enzyme that palmitoylates DLK in neurons, and loss of ZDHHC17 protects against degeneration (Niu *et al*, 2022). Whether DLK is involved in p75NTR retrograde signaling remains an interesting question for future studies.

These parallel findings raise the possibility that palmitoylation is a common enabling modification for long-distance regressive signaling modules, in marked contrast to progressive signaling endosomes carrying TrkA, which do not require palmitoylation for retrograde movement (Ascano *et al*, 2012). Whether other regressive signaling receptors, such as PlexinA3 or TNFR superfamily members that undergo regulated intramembrane proteolysis, employ similar mechanisms remains to be tested. Identifying the relevant DHHC enzyme(s) for p75NTR is a priority, as this would test whether a single enzyme coordinates multiple regressive modules or whether there exists a “palmitoylation code” across different death pathways.

### Implications for Functionally Distinct Pools of Signaling Endosomes

Our findings speak to a central unresolved question in the long-distance signaling field: are there functionally distinct “flavors” of signaling endosomes, or does a single population of vesicles carry multiple signals whose interpretation depends on context? The observation that p75NTR-positive endosomes are marked by CD63 and travel retrogradely in a RAB5- and RAB7-dependent manner (Pathak *et al*, 2018; Escudero *et al*, 2019) overlaps substantially with the trafficking itinerary described for TrkA-positive progressive endosomes (Deinhardt *et al*, 2006). Yet the functional consequences, survival versus death, are diametrically opposed.

If palmitoylation status determines which receptors can be loaded into particular endosomal carriers, then lipid modifications could provide a sorting mechanism that segregates progressive and regressive signals into distinct vesicular pools despite their use of common trafficking machinery. Alternatively, palmitoylation might enable p75ICD to associate with a subset of endosomes containing regressive signaling components such as activated JNK3, DLK, or specific adaptor proteins, thereby creating functionally specialized carriers even within molecularly overlapping populations. Testing these models will require approaches such as super-resolution co-localization of TrkA or DLK with p75NTR (WT vs. Δpalm) on the same endosomes, or proteomic comparison of endosomes isolated under survival versus death conditions.

### AND-Gate Architecture: Palmitoylation Plus Proteolysis

Our data demonstrate that palmitoylation and cleavage are functionally separable: the p75NTR^Δpalm^ mutant undergoes normal proteolytic processing (Fig. 5) but fails to internalize or signal death, while prior work showed that inhibition of γ-secretase prevents death even when palmitoylation can proceed normally (Kenchappa *et al*, 2006; Pathak *et al*, 2018). This independence suggests that palmitoylation and proteolysis represent distinct checkpoints that must both be satisfied for productive regressive signaling. Moreover, it provides an elegant mechanism for fine-tuned regulation and spatiotemporal control of p75NTR degenerative signaling.

### Long-Distance Regressive Signaling in Development and Disease

The broader implications of our findings extend to long-distance regressive signaling in neural development and pathology (Pathak *et al*, 2021). During nervous system assembly, long-distance death cues help sculpt circuits by eliminating neurons that innervate inappropriate targets or fail to secure adequate trophic support (Rabizadeh *et al*, 1993; Oppenheim, 1991). Our identification of palmitoylation as a key enabler of p75NTR retrograde death signaling adds to an emerging picture that includes requirements for p150^Glued^ deacetylation by axonal HDAC1 (Pathak *et al*, 2018)) and JNK activation in distal axons (Escudero *et al*, 2019).

The re-engagement of developmental death pathways in neurodegenerative disease is well established. In amyotrophic lateral sclerosis (ALS), motor neurons display increased retrograde transport of pro-degenerative signals coincident with decreased transport of pro-survival signals (Perlson et al., 2009). In Alzheimer’s disease, p75NTR binds amyloid-β peptides and may facilitate their effects on basal forebrain cholinergic neurons (BFCNs) (Hu *et al*, 2013; Ovsepian *et al*, 2014). Friedman and colleagues reported that BFCNs undergo retrograde degenerative signaling following a concussive impact injury at their terminals in the cortex (Dasgupta *et al*, 2023). The injury induced the local production of pro-neurotrophins and the neuronal death back in the basal forebrain was prevented by deletion of p75NTR. Similarly, this same group demonstrated that addition of proNGF to distal axons of BFCNs in compartmentalized cultures induced p75NTR-dependent neuronal apoptosis, similar to our previous findings in sympathetic neurons (Pathak *et al*, 2018) and Fig. 6). Interestingly, this retrograde signal also involved local translation of amyloid precursor protein, which was necessary for the subsequent apoptosis (Dasgupta *et al*, 2024). Although our current data are restricted to developmental sympathetic neurons, the parallels with known disease and injury pathways suggest that dysregulated palmitoylation of regressive signaling modules is an attractive hypothesis to test in future studies of neurodegeneration.

### Outstanding Questions and Future Directions

Our work establishes palmitoylation as essential for p75NTR endocytosis and long-distance regressive signaling but raises key mechanistic questions. Which DHHC palmitoyl acyltransferase(s) modify p75NTR, and where and when are they active? Which acyl-protein thioesterases remove palmitate from p75NTR, and could dynamic palmitoylation cycles regulate receptor function? Does palmitoylation influence p75NTR interactions with specific co-receptors such as Sortilin or Lingo-1, or adaptors such as NRIF? Testing whether other regressive signaling receptors require palmitoylation for long-distance signaling could reveal unifying principles for how the nervous system propagates death information over long distances.

## Methods

### Rats and Mice

All animal experiments were approved by the Institutional Animal Care and Use Committee at Vanderbilt University School of Medicine and the University of Virginia. Wild type, untimed pregnant-CD1 rats and wildtype-C57BL/6J mice were purchased from Charles River Laboratory, USA. P75NTR^Δpalm/Δpalm^ mutant mice were produced in the C57BL/6J background at the Genetically Engineered Murine Model (GEMM) core, University of Virginia. Wildtype-C57BL/6J, P75NTR^Δpalm^, or C57BL/6Jp75NTR^(exonIII)-/-^ mice and pregnant rats were housed in 12-hour day and night cycle environment with ad libitum availability of chow diet and water. Both male and female postnatal rats or mice of postnatal day 0-3 age were used in all the experiments, as indicated.

### Generation of P75NTR^Δpalm/Δpalm^ mice by CRISPR mutagenesis

We used CRISPR/Cas9-mediated genome editing with the help from Wenhao Xu at the Genetically Engineered Murine Model (GEMM) core, University of Virginia. Using a CRISPR repair template strategy Cysteine 281 (analogous to rat C279) was replaced with alanine, rendering p75NTR receptor palmitoylation-resistant in the p75NTR^Δpalm/Δpalm^ mutant mice.

The nucleotides TG were changed to GC in exon 5 of *ngfr* gene on chromosome 11 to replace the TGC codon in the amino acid chain for cysteine 281 with GCC (alanine) to generate non palmitoylable, C281A mutant p75NTR. This mutation also created a novel PvuII restriction digestion site at 5′ of the knock-in C281A replacement codon in *ngfr* which allows straightforward genotyping of the mice by PvuII digestion following PCR amplification. In addition to the mutation at C281, we also introduced eight more downstream silent mutations in the repair template oligonucleotide to prevent nonspecific digestion by CAS9. The eight silent mutations were introduced immediately next to C281 and are present in the mutant allele but not in WT. The CAS 9 nuclease, a guide RNA with the oligonucleotide for repair template carrying the desired mutations and homology arms, were co-injected into the male pronucleus of fertilized eggs, followed by implantation of the eggs into surrogate mothers to obtain offspring. We developed an allele-specific primer for genotyping the pups and obtained two male and two female P75NTR^Δpalm/Δpalm^ founder mice that appeared to be heterozygous for the mutation (1:1 peak ratio from sequencing). From F1 pups generated from the founder mice, we bred heterozygous males and females to obtain homozygous P75NTR^Δpalm/Δpalm^ mutant mice. The homozygous mice are viable and have been backcrossed to WT to obtain congenic animals (7 generations of backcrossing), thereby removing unlinked off-target mutations. The palmitoylation resistant p75NTR^Δpalm/Δpalm^ mice expressed similar levels of p75NTR but it cannot be palmitoylated as confirmed by Acyl RAC assay and immunoblotting with anti-p75NTR antibody (Fig. 7B & 7C).

### Cell Culture and Treatments

Sympathetic neurons were isolated from superior cervical ganglia, dissected from male and female postnatal day 0-3 old mice or rats and dissociated with 0.08% trypsin (Worthington), 0.3% collagenase and DNAse (Sigma). Dissociated cells were then plated on poly-d-lysine (MP Biomedicals) and laminin (Invitrogen) coated surface. For mass cultures, neurons were cultured in UltraCULTURE medium (Lonza) supplemented with 3% fetal bovine serum (Invitrogen), 20 ng/ml nerve growth factor (Harlan), 2 mM L-glutamine (Invitrogen), 100 units/ml penicillin (Invitrogen), and 100 µg/ml streptomycin (Invitrogen) or as indicated. 24 hours after plating, the neurons were treated with 10 µM cytosine arabinofuranoside (AraC) (Sigma) for 24 hrs to inhibit the proliferation of non-neuronal cells. Following 2 days of exposure, AraC was removed and after 24 hrs, the neurons were treated at the indicated concentrations with the reagents as described in respective figure legends. Due to lack of availability of UltraCULTURE media, as Lonza ceased production of the UltraCULTURE media, the neurons used for experiments assessing p75NTR internalization were cultured in DMEM/F12 (Gibco) supplemented with PRIME-XV IS21 (Fujifilm Biosciences). Neurons cultured using IS21X supplement were not treated with AraC. For all experiments involving NGF removal and treatment with 200ng/ml BDNF in presence or absence of 2-bromo palmitate (Sigma) at a concentration of 100µM, the neurons were rinsed at least 3 times with media lacking NGF, containing 12.5 mM KCl and 100 ng/ml of anti-NGF antibody with at least 10 min incubation each in between the rinses prior to addition of the indicated reagents. The neurons were treated for time durations as indicated in the figure legends.

For neuron cultures in microfluidic devices (from Xona Microfluidics or generated in-house in the laboratory of Christopher Deppmann, University of Virginia), the chambers were sterilized in plasma cleaner and assembled on coverslips, then coated with poly D lysine and laminin as indicated above for neuron mass cultures. Dissociated neurons were plated on one side (proximal) of the devices in ultraCULTURE media or DMEM/F12 with 20 ng/ml NGF and axons were allowed to grow and cross into the other side (distal). To direct the growth of axons towards the distal side, media with 50 ng/ml NGF was added in the distal compartment for the initial two days. We confirmed that there was no diffusion of soluble material between the chambers using a fluorescent dye (Cy3). After the axons reached the distal side, the neurons were kept alive by maintaining the cell bodies in media containing 20 ng/ml NGF or 12.5 mM KCl and distal axons were treated with BDNF in presence or absence of 100 µM 2-Bromo palmitate as indicated in figure legends. After the treatments, neurons were fixed in 4% PFA for immunostaining and counting axonal puncta or apoptotic nuclei.

COS7 cells and HEK293 cells were cultured in high glucose DMEM with 10% fetal bovine serum and penicillin /streptomycin (100 U/ml, 100 μg/ml, respectively) and transfected using Lipofectamine 2000 (Invitrogen), according to the manufacturer’s instructions.

### Plasmid DNA constructs

Wildtype full length rat p75NTR-GFP expression construct, expressing p75NTR with GFP fused to the C-terminus, was kindly provided by Dr. Eran Perlson (Tel Aviv University, Israel). The WT rat p75NTR-GFP construct on eGFP reporter cassette (Addgene plasmid Plasmid #11795) was mutated to palmitoylation resistant C279A p75NTR^Δpalm^ construct wherein we mutated cysteine 279 residue to alanine by site directed mutagenesis. Wildtype full length HA-p75NTR and wildtype p75ICD only rat constructs from the Carter lab (Kenchappa *et al*, 2006) were also mutated to respective palmitoylation resistant C279A forms by site directed mutagenesis. The accuracy of nucleotides in all the wild type constructs and the changes introduced in the mutant constructs were confirmed by sequencing the wild type and mutant constructs.

### Lentivirus production and infection

Lentiviral particles for p75NTR-GFP fusion protein expression in primary sympathetic neurons were prepared by transfecting WT-p75NTR-GFP or non-palmitoylable C279A mutant P75NTR^Δpalm^ expression plasmids, along with the packaging vectors, pSPAX2 and pMD2.G (Addgene plasmids 12260 and 12259, respectively) using polyethylenimine (Polysciences) into LentiX-HEK293T cells (Takara Bio USA, Inc., Cat# 632180) grown in 15-cm Petri dishes to ∼80% confluence in DMEM (Gibco) supplemented with 10% fetal bovine serum (Gibco). We did each transfection with 15 ug total DNA at 50:37.5:12.5% ratio of expression vector:pSPAX2:pMD2.G respectively followed by lentivirus conditioned medium collection after 24 and 48 hrs. We concentrated the lentiviral particles in a Beckman Coulter Optima LE-80K Ultracentrifuge for 2 h at 23,000 rpm (95,000 g) at 4°C in an SW28 rotor, resuspended them in 1 ml DMEM medium and stored at -80°C in 20 ul aliquots. Cultured neurons were transduced in UltraCULTURE medium or DMEM/F12 media with a viral MOI of 5 and incubated for 48-72 h before assays were performed.

### Detection of palmitoylated proteins by Acyl-RAC assay

COS7 cells or neurons expressing full length (p75NTR) or p75ICD fragments were collected in lysis buffer (100mM HEPES, 1 mM EDTA, 2.5% SDS, protease inhibitor cocktail, pH7.5) for sonication in cuphorn sonicator at room temperature (RT). To block free thiols, we used 1% methyl methane thiosulfonate (MMTS, pH 7.5) and incubated the lysates at 40°C for 10 minutes with frequent intermittent vortexing. After addition of MMTS, we saved approximately 5% (20ul) of each supernatant as “input.” Lysates were centrifuged at 13,000 rpm for 10 minutes at RT in a microfuge and supernatants were transferred to another tube before adding three volumes of 100% acetone followed by incubation at -20C to precipitate the proteins. After 20 minutes, the solution was centrifuged at 7300 rpm for 5 minutes to pellet the precipitated proteins. The pellets were washed five times with prechilled 70% acetone at 7,000 rpm for 1-2 minutes each. After the final wash, pellets were air dried to remove residual acetone. We resuspended pellets in 600ul binding buffer (100mM HEPES, 1 mM EDTA, 1.0% SDS, pH7.5) and sonicated for 5-10 seconds, if required, to completely dissolve the pellets. After complete resuspension, the solution was divided into two tubes and either 2M hydroxylamine (HAM) or 2M NaCl added. Then pre-equilibrated thiopropyl (TP)-sepharose beads (about 50µl settled bead volume) were added to each tube and shaken for 2 to 4 hours at RT to bind the palmitoylated protein. Covalently linked TP sepharose-protein beads were washed five times in binding buffer (100 mM HEPES 1 mM EDTA and 1.0% SDS). For immunoblot analysis, palmitoylated proteins were eluted in 50ul of 2X Laemmeli buffer with DTT. Samples with 2M NaCl served as negative (-HAM) control for palmitoylation assay.

### Immunostaining

Primary wildtype sympathetic neurons cultured in microfluidic devices were treated on their distal axons or cell bodies as indicated then fixed with 4% paraformaldehyde (PFA). Fixed neurons were washed with PBS and blocked with 5% normal donkey serum in 0.1% TritonX 100 in PBS for 1 hr at RT followed by incubation with primary antibodies rabbit anti-p75ICD antibody (Abcam #ab52987) and mouse TUJ1 at 1:500 diluted in blocking solution. Similarly, p75-/- neurons expressing full length WT-p75NTR-GFP or C279A-mutant p75NTR^Δpalm^-GFP were stained with mouse anti-GFP antibody (Clontech, 632380) at 1: 500 dilution. After overnight incubation in primary antibody solution, they were rinsed and incubated with fluorescently tagged secondary antibodies, Goat anti-rabbit-488 (Invitrogen cat# A11008) and Donkey anti-mouse-568 (Invitrogen cat# A10037). Nuclei were visualized by DAPI, and images were acquired with a confocal laser imaging system (LSM 980; Carl Zeiss MicroImaging, Inc.). Endogenous p75ICD stained vesicles in the axons labeled with TUJ1 were counted blindly and 30-40 axons were scored per condition in each experiment.

To quantify p75NTR internalization, primary sympathetic neurons from WT and p75NTR^Δpalm^ mice were grown in 20 ng/ml NGF containing media. After three DIV, NGF was removed by replacing the media with DMEM containing 1 mg/ml BSA and 12.5 mM KCl for serum deprivation. After 1 hour, anti-p75ECD antibody with ATTO-550 at 1:100 (Alomone labs) was added for 2 hrs at 4°C. After removing the anti-p75NTR antibody, neurons were left in KCl or treated with 200 ng/ml BDNF at 37°C for 30 minutes to several hours as indicated in the figure legend. The labelled neurons were imaged on Zeiss LSM 710 confocal microscope with a Plan apochromat 40X/1.4 water differential interference contrast (DIC) objective. Z-stack images were taken to capture the total intensity associated with the cell, including the interior and the cell surface. Brightfield images were taken to identify cell membranes. In all the images, the same number of z-slices were collapsed in Zen black for uniform thickness across the cells. Then cell-associated fluorescence was quantified in Zen blue with internalized p75NTR being the receptor present in the cell interior and p75NTR present at the cell surface was determined by subtraction of internalized p75NTR from the total intensity associated with the cell.

To assess the ligand induced localization of p75NTR in acidic vesicles, p75^-/-^ neurons expressing full length WT-p75NTR-GFP or C279A-mutant p75NTR^Δpalm^-GFP were stained live with pHrodo labeled anti-p75NTR antibody after serum deprivation. Briefly, Anti-p75NTR antibody (Alomone Cat# ANT-007) was labeled with pHrodo iFL Red following manufacturer’s instruction (invitrogen). pHrodo iFL Red was dissolved in DMSO to prepare the 2 mg/mL (2 mM) pHrodo™ iFL labeling solution and added to BSA free Anti-p75NTR antibody (1mg/ml). The labeled antibody was purified by the gel resin spin column provided in the pHrodo™ iFL Red Microscale Protein Labeling Kit. After an hour of serum deprivation of whole neurons in DMEM/F12 media with 1mg/ml BSA, distal axons were exclusively treated with BDNF or KCl along with pHrodo-p75NTR antibody as indicated and imaged live on Zeiss LSM980.

### Biotinylation assay

The neurons cultured from rat SCGs were infected with lentivirus to express either WT- or p75NTR^Δpalm^-GFP. To begin with the biotinylation assay, neurons were cooled to 4°C by placing the dishes on an ice bath in the cold room and rinsed twice with ice cold PBS containing 1.5 mM MgCl2, 0.2 mM CaCl2 (PBS^2+^). Surface proteins were biotinylated by adding freshly prepared 10mM Sulfo-NHS-SS-Biotin and incubation at RT for 30 minutes. To quench the biotin reaction, ice-cold quenching solution (PBS^2+^ supplemented with 100 mM glycine) was added for 5 minutes on ice after removing the biotin solution, then rinsed twice in ice cold quenching solution for fifteen minutes each at 4°C. The biotinylated surface proteins were stripped with freshly prepared 50 mM of Tris(2-Carboxyethyl) phosphine Hydrochloride (TCEP) by incubating them twice in ice cold stripping buffer for fifteen minutes each at 4°C. The cells were then rinsed 3 times with neutralization buffer (150 mM NaCl, 1.0 mM EDTA, 0.2% BSA, 20 mM Tris, pH 8.6), at 4°C followed by washing with PBS^2+^ before preparing the lysates. The neurons were lysed in RIPA buffer containing fresh protease inhibitors (1.0 mM PMSF, 1.0 μg/ml each leupeptin, aprotinin and pepstatin). Protein concentration was estimated by BCA assay and then streptavidin affinity chromatography was done by adding a suspension of streptavidin agarose beads at 20μl beads/50 μg lysate and incubated overnight at 4°C on a tube rotator. The biotin bound proteins were then eluted in reducing Laemmli sample buffer after gently washing the beads with RIPA buffer. The eluates were run on a Tricine gel followed by immunoblotting with anti-GFP antibody to detect p75NTR-GFP.

### Immunoblotting

For p75NTR immunoblotting, whole cell lysates from wildtype or mutant overexpressing cells were prepared in 2X Laemmli buffer after rinsing with ice-cold PBS. Cell lysates were subjected to Tricine-Tris SDS-PAGE, transferred to nitrocellulose, blocked in 10% skimmed milk and probed using an antibody against p75NTR ICD (Kenchappa *et al*, 2006) at a dilution of 1:1000.

### Genotyping P75NTR^Δpalm/Δpalm^ C281A mutant mice

DNA isolated from mice ear punches was amplified using high fidelity PCR reaction to amplify the target region containing the novel PvuII restriction digestion site generated at 5′ of the knock-in C281A replacement codon. The PCR product was digested by PvuII and electrophoresed on agarose gel to identify the WT or P75NTR^Δpalm/Δpalm^ C281A mutant mice based on the digestion of PCR amplicons. The genotypes were also confirmed by Sanger sequencing of the PCR amplicons.

### Sectioning and Immunostaining of Superior Cervical Ganglia (SCG)

SCGs from WT, p75^Δpalm/Δpalm^, p75^-/-^ mice at postnatal day 0 were isolated and fixed in 4% PFA for 24 hours. The ganglia were then cryopreserved in 30% sucrose, embedded in OCT media and the entire ganglia was cryosectioned at 20 µM thickness. The sections were subjected to antigen retrieval in sodium citrate buffer (10mM, pH 6.0) for 20 mins at 65°C, washed in PBS following permeabilization and blocking in 0.2% Triton X-100, 5% normal donkey serum for 1 hour at room temperature (RT). Primary antibodies against cleaved caspase 3 (Cell signaling technology, 9661) and Tuj1 (Covance, MMS-435-P) were applied at 1:250 dilution in blocking buffer overnight at 4°C. After removal of primary antibody sections were rinsed in PBS and subtype-specific Alexa Fluor fluorescent secondary antibodies were added for 1 hour at RT. The sections were rinsed in PBS and mounted with ProLong Gold antifade reagent with 4’,6-diamidino-2-phenylindole dihydrochloride (DAPI).

For quantitative assessment of apoptotic neurons, images of the SCG sections were captured using a Nikon Eclipse Ti fluorescence microscope. The cleaved caspase 3 and Tuj1 double positive cells were quantified in every other section and normalized to the total number of cells in the section (determined using DAPI and Fiji software) through the entire ganglion.

## Acknowledgements

“This work used genetically engineered P75NTR^Δpalm/Δpalm^ mouse strain which were produced at the Genetically Engineered Murine Model Core Facility which is supported by the University of Virginia School of Medicine, Research Resource Identifiers (RRID):SCR_025473. This study was supported by DBT RRF (BT/HRD/35/02/2006) to AP and NIH R01 NS136342 to C.D.D. and B.D.C.

